# Comprehensive interrogation of human skeletal muscle reveals a dissociation between insulin resistance and mitochondrial capacity

**DOI:** 10.1101/2023.02.24.529750

**Authors:** KL Whytock, MF Pino, Y Sun, G Yu, FG De Carvalho, RX Yeo, RB Vega, G Parmar, A Divoux, N Kapoor, F Yi, H Cornnell, DA Patten, ME Harper, SJ Gardell, SR Smith, M Walsh, LM Sparks

**Author notes:** Corresponding Author: 301 E Princeton St, Translational Research Institute, AdventHealth, Orlando, FL, 32804.

## Abstract

**Aims/Hypothesis:** Insulin resistance and blunted mitochondrial capacity in skeletal muscle are often synonymous; however, this association remains controversial. The aim of this study was to perform an in-depth multi-factorial comparison of skeletal muscle mitochondrial capacity between individuals who were lean and active (Active), individuals with obesity (Obese) and individuals with Obesity, insulin resistance and type 2 diabetes (T2D).

**Methods:** Skeletal muscle biopsies were obtained from the *Vastus Lateralis* of individuals who were lean and active (Active- n = 9), individuals with obesity (Obese- n = 9) and individuals with obesity insulin resistance and T2D (T2D- n =22) in this cross-sectional design. Mitochondrial capacity was assessed by *ex vivo* mitochondrial respiration with fatty-acid and glycolytic supported protocols adjusted for mitochondrial content (mtDNA and citrate synthase activity). Supercomplex assembly was measured by BN-PAGE and immunoblot. TCA cycle intermediates were assessed with targeted metabolomics. Exploratory transcriptomics and DNA methylation analyses were performed to uncover molecular differences affecting mitochondrial function among the three groups.

**Results:** Active had greater mitochondrial capacity compared to both Obese and T2D for *ex vivo* mitochondrial respiration with fatty-acid and glycolytic supported protocols adjusted for mitochondrial content (*P* < 0.05). Complex IV supercomplex assembley was greater in Active compared to Obese and T2D (*P* < 0.05) whereas Complex I and III supercomplex assembly was greater in Active compared to T2D only (*P* < 0.05). TCA cycle intermediates; citrate, succinate, fumarate and malate were all significantly greater in Active compared to Obese and T2D (*P* < 0.05). Strikingly, Obese and T2D do not differ in any of the skeletal muscle mitochondrial measurements. Active had an upregulation of genes related to respiration/mitochondrial capacity compared to both Obese and T2D. Transcriptional differences between Obese and T2D were not driven by mitochondrial related process. Active had reduced methylation correlated with increased gene expression for important mitochondrial-related genes, including *ATP5PD* and *MFN2*.

**Conclusions/Interpretations:** We reveal no discernable differences in skeletal muscle mitochondrial content, mitochondrial capacity and mitochondrial molecular profiles between obese individuals with and without T2D that had comparable levels of confounding factors (BMI, age, aerobic capacity) that affect mitochondrial capacity. We highlight that lean, active individuals have enhanced skeletal muscle mitochondrial capacity that is also reflected at the level of DNA methylation and gene transcription. The collective observation of comparable muscle mitochondrial capacity in individuals with obesity and T2D (vs. individuals without T2D) underscores a dissociation from skeletal muscle insulin resistance.

**Clinical trial number:** NCT0191110

## Introduction

Skeletal muscle (SkM) insulin resistance is a primary defect in the pathology of type 2 diabetes (T2D) [1,2]. Insulin resistance in SkM has historically been characterized by impaired mitochondrial oxidative capacity [3]. Dysregulated oxidation of lipid species is proposed to contribute to lipotoxicity in SkM and associated with insulin resistance [4,5]. However, the apparent association between SkM insulin resistance and impaired mitochondrial capacity is not always evident. Endurance exercise training in individuals with T2D simultaneously improves insulin sensitivity and SkM mitochondrial capacity quantified *in vivo* as phosphocreatine (PCr) recovery rate [6,7] and *ex vivo* by high resolution respirometry in permeabilized SkM fibers [8]. In contrast, thiazolidnediones improves insulin sensitivity in individuals with T2D without impacting SkM mitochondrial capacity quantified *in vivo* with PCr recovery rate [9,10].

At a cross-sectional level, individuals with T2D have comparable *in vivo* SkM mitochondrial capacity (quantified as ATPmax) to obese controls [11]. Additionally, at least half of a cohort of individuals with T2D showed comparable ATPmax to sedentary non-obese controls [11], highlighting the heterogeneity in the pathophysiology of T2D. SkM from individuals with obesity and with or without T2D have comparably lower succinate dehydrogenase enzyme activities (a spectrophotometric measure of mitochondrial activity in SkM tissue) compared to lean individuals [12]. In contrast, oxidative pathway enzyme activities (citrate synthase, cytochrome-c oxidase) are lower in SkM from individuals with T2D compared to lean and obese non-diabetic individuals [13]. Furthermore, isolated mitochondria from T2D SkM showed modest reductions in *ex vivo* respiration normalized to citrate synthase activity compared to healthy individuals with obesity [14]. While another report [3] found a hierarchical order of impaired capacity of the respiratory chain quantified by NADH:O2 oxidoreductase activity in SkM from T2D, obese and lean individuals. Discrepancies among these results may partially be attributed to the techniques employed for measurements of SkM mitochondrial capacities (enzymatic activities, *ex vivo* respiration, *in vivo* ATP max/PCr recovery rates, transcriptomics) and phenotypic differences (aerobic capacity, BMI, age) among the groups.

In the study we leveraged previously deeply phenotyped samples from a previous study to interrogate differences in mitochondrial function in individuals who were lean and active (Active), individuals with obesity (Obese) and individuals with Obesity, insulin resistance and type 2 diabetes (T2D) [15]. Importantly, in this cohort Obese and T2D were matched for age, BMI and aerobic capacity. The aim of the study was to perform an in-depth multi-factorial assessment of mitochondrial function using; *ex vivo* respiration adjusted for mitochondrial content, supercomplex assembly and TCA cycle intermediates. In parallel, we performed exploratory multi-omics (transcriptomics and DNA methylation) of SkM to uncover molecular differences affecting mitochondrial function.

## Research Design and Methods

### Human Participants

The samples and acquired data used in this study were part of a larger clinical trial comparing variation in endurance exercise response [6] under clinical trial number NCT01911104. We selected data and samples from 22 individuals with T2D (T2D, Males = 13), 9 participants with obesity (Obese, Males = 2) and 9 Active participants (Active, Males = 9) for this cross-sectional interrogation. Phenotypic differences of this cohort, including aerobic capacity measured by VO2 peak, body composition by DXA, *in vivo* mitochondrial capacity measured by PCr recovery rate and insulin sensitivity using hyperinsulinemic euglycemic clamp have previously been published [15]. This data is included in participant characteristics table 1 for reference. The study protocol was approved by the AdventHealth Institutional Review Board and carried out in accordance with the Declaration of Helsinki. All participants provided written informed consent. At the time of enrollment individuals with T2D had to have an HbA1c ≤ 8% if receiving glucose-lowering medication and between 6 and 8.5% if treated with diet alone. Participants ceased glucose-lowering treatment two weeks prior to participation. All measurements were conducted following an overnight fast.

**Table 1.** Clinical characteristics.

| Clinical characteristics | T2D<br>(n= 22) | Obese<br>(n=9) | Active<br>(n=9) |
| --- | --- | --- | --- |
| Age (years) | 50.1 ± 8.1 | 46 ± 8.8<sup>#</sup> | 33.7 ± 9.3<sup>*</sup> |
| Biological sex (F/M) | 9/13 | 7/2 | 0/9 |
| BMI (kg/m<sup>2</sup>) | 36.8 ± 5.6 | 38.5 ± 8.5<sup>#</sup> | 23.4 ± 1.9<sup>*</sup> |
| Body Mass (kg) | 106.0 ± 20.3 | 100.9 ± 27.2<sup>#</sup> | 77.5 ± 9.9<sup>*</sup> |
| Fat Mass (kg) | 44.7 ± 14.1 | 47.3 ± 16.2<sup>#</sup> | 16.2 ± 6.4<sup>*</sup> |
| Fat-Free Mass (kg) | 61.2 ± 11.1 | 53.6 ± 13.9 | 61.4 ± 6.7 |
| HbA1c (%) | 7.3 ± 0.9<sup>&</sup> | 5.6 ± 0.4 | 5.2 ± 0.4<sup>*</sup> |
| T2D duration (years) | 4.6 ± 4.7 | - | - |
| M Value (μmol/kgFFM/min) | 5.0 ± 2.8<sup>&(P = 0.05)</sup> | 9.4 ± 5.5 | 13.0 ± 5.0<sup>*</sup> |
| Fasting glucose (mg/dl) | 182.2 ± 56.3<sup>&</sup> | 94.4 ± 11.1 | 89.8 ± 4.9<sup>*</sup> |
| Fasting insulin (μIU/ml) | 16.4 ± 7.3<sup>&</sup> | 10.3 ± 3.9<sup>#</sup> | 2.6 ± 0.9<sup>*</sup> |
| Free fatty acids (mmol/l) | 0.5 ± 0.2 | 0.5 ± 0.2 | 0.3 ± 0.1<sup>*</sup> |
| Total cholesterol (mg/dl) | 152.8 ± 21.1 | 162.9 ± 35.6 | 156.4 ± 24.3 |
| LDL (mg/dl) | 90.2 ± 23.6 | 96.6 ± 24.5 | 90.2 ± 19.1 |
| HDL (mg/dl) | 37.7 ± 8.5 | 44.5 ± 7.9 | 49.8 ± 7.7<sup>*</sup> |
| Triglycerides (mg/dl) | 125.8 ± 40.6 | 108.9 ± 86.9 | 81.9 ± 29.8 |
| PCr recovery rate (1/seconds) | 0.020 ± 0.007 | 0.023 ± 0.010<sup>#</sup> | 0.041 ± 0.010<sup>*</sup> |
| VO<sub>2 peak</sub> (L/min) | 2.29 ± 0.68 | 2.24 ± 0.86<sup>#</sup> | 4.17 ± 0.64<sup>*</sup> |
| VO<sub>2 peak</sub> (ml/kg/min) | 22.7 ± 5.6 | 22.1 ± 6.1<sup>#</sup> | 54.3 ± 5.2<sup>*</sup> |
| VO<sub>2 peak</sub> (ml/kg FFM/min) | 38.1 ± 5.3 | 40.5 ± 8.1<sup>#</sup> | 67.8 ± 6.6<sup>*</sup> |

Data are the mean ± SD. F, female; M, male. T2D, type 2 diabetic. Obese, obese non-T2D. Active, lean Active. BMI, Body mass index. FFM, Fat-free mass. M-value, whole-body insulin sensitivity.
HbA1c, hemoglobin A1C. VO<sub>2peak</sub>, maximum rate of oxygen consumption. LDL-c, low density lipoprotein. HDL-c, high density lipoprotein. PCr, Phosphocreatine. VO<sub>2peak</sub>, maximum rate of oxygen consumption. <sup>\*</sup>difference between Active and T2D. <sup>&</sup>difference between T2D and Obese.
<sup>#</sup>difference between Active and Obese. One-way ANOVA, *P* <0.05.

### Skeletal Muscle biopsy

SkM biopsies were performed in the *vastus lateralis* using Bergström needle technique [16] under fasting conditions. Portions of SkM (150 mg) were snap frozen for Supercomplex assembly quantification, targeted metabolomics, RNA and DNA extractions, and a fresh portion (10 mg) was used for assessment of mitochondrial respiration. Certain assays were restricted to SkM availability.

### Citrate Synthase Activity

Citrate synthase activity in SkM tissue was assessed spectrophotometrically. Assay buffer consisting of Buffer Z (https://tinyurl.com/2kuthy7r (no BSA)) supplemented with 5’,5’-Dithiobis 2-bitrobenzoic acid and acetyl-CoA was dispensed with permeabilizing agent Alamethicin and SkM lysate and incubated at 37°C for 5 min. The reaction was initiated by oxaloacetate. Absorbance was recorded at 412 nm every 1 minute for 20 cycles. Citrate synthases activity was calculated using the Beer-Lambert Law with a molar absorption coefficient of thionitrobenzoic acid (13.6 mM/cm).

### mtDNA copy number

Total DNA was isolated from 10-20 mg of SkM tissue using DNeasy blood and tissue extraction kit (QIAGEN Inc, Valencia, CA). Relative amounts of mitochondrial DNA (mtDNA) and nuclear DNA were quantified by real-time quantitative PCR [17].

### High-resolution respirometry

Mitochondrial respiration on isolated permeabilized SkM fibers using high-resolution respirometry with a pyruvate simulated glycolytic protocol and a fatty acid protocol has previously been reported in this cohort [15]. We leveraged this previously published data to assess whether differences in mitochondrial capacity were due to function or content of mitochondria by adjusting these respiration values to mitochondrial content measured by mtDNA and citrate synthase activity. For full details of the respirometry protocol the reader is directed to Carnero et al. [15]. During each protocol, respiration of individual complexes were quantified; Complex I supported LEAK (L_I_ or (L_FAO_), complex I supported oxidative phosphorylation (OXPHOS; P_I_ or P_I +FAO_), Complex I and II supported OXPHOS (P_I+II_ or P_I+II +FAO_) maximal electron transfer system capacity (E_I+II_ or E_I+II +FAO_).

### Supercomplex immunoblots

Electron transport system (ETS) supercomplex (SC) assembly was analyzed by blue native polyacrylamide gel electrophoresis and blotting of individual components with immunoblotting [18–20]. SC and monomers were normalized to nuclear-encoded Complex II which does not participate in mammalian ETS SC formation [19].

### Tricarboxylic cycle intermediates

Tricarboxylic (TCA) cycle intermediates (TCAi) were measured by targeted LC-MS/MS [21]. Samples were spiked with heavy isotope-labelled internal standards and derivatized with O-benzylhydroxylamine (OBA) and 1-ethyl-3-(3-dimethylaminopropyl) carbodiimide before being separated on a Waters Acquity UPLC BEH. Quantitation of OAs was performed by single reaction monitoring using a Thermo Scientific Quantiva triple quadrupole mass spectrometer (Thermo Scientific, San Jose, CA). The raw data was processed using Xcalibur 3.0.

### RNA sequencing

RNA-Seq on SkM tissue was performed on a subset of participants from each group (n = 6, per group). RNA was extracted from 30-50mg of SkM tissue using RNeasy Fibrous Tissue Kit (Qiagen, Valencia, CA). Following polyA selection, library preparation and quality control, sample libraries underwent mRNA sequencing, with the average depth of 20 million paired-end reads per sample using the Illumina Novaseq 6000. Adaptor was trimmed with Cutadapt and low-quality reads were filtered out before aligning with STAR to the reference genome hg38. Raw counts were calculated with featureCounts. Cufflinks was used to generate normalized read counts per gene and isoform in terms of FPKM values.

### Reduced representation bisulfite sequencing

RRBS (Reduced representation bisulfite sequencing) on SkM tissue was performed on a subset of participants from each group (n = 3, per group). Total DNA was isolated from 10-20 mg of SkM tissue using DNeasy blood and tissue extraction kit (QIAGEN Inc, Valencia, CA). RRBS library was generated from ≥ 1.5 μg genomic DNA at Novogene. Libraries were sequenced as Paired-end 150bp on Novaseq 6000 sequencer at an average out per sample ≥ 10 Gb.

### Quantification and statistical analyses

To compare differences between groups for clinical characteristics and measures of mitochondrial capacity/content an ANOVA model with Tukey post hoc adjustment was applied and significance was set at *P* < 0.05. Respirometry data adjusted for mitochondrial content (citrate synthase activity or mtDNA copy number) were analyzed using an ANCOVA followed by post-hoc comparisons with significance set at *P* < 0.05. Statistical analyses were performed in JMP.

### Bioinformatics

#### Transcriptomics

The data were first log_2_-transformed and then z-scaled for transformation and standardization respectively. Principal component analysis (PCA) was performed with factoextra and ggplot2 using top 1624 genes ranked by Randomforest classification [22]. Differential expressed genes (DEG) were analyzed with limma R package for between group differences [23] and adjusted for sex, age and BMI. Gene-ontology (GO) over-representation analysis on DEGs (P<0.05) was conducted using clusterProfiler, using genes detected as a background list and an FDR of <0.005 [24]. Redundancy of GO terms from DEGs was reduced with GOSemSim [25].

#### DNA Methylation

Initially, Trim Galore, a wrapper tool around Cutadapt and FastQC, was applied to FastQ files for quality control and adapter trimming. Once trimmed, Bismark was performed for read mapping and methylation calling [26]. Cytosines in the reference sequenced as thymines are labeled unmethylated, and those that remained as cytosines are labeled methylated. Differentially methylated site (DMS) analysis was performed with R package methylKit [27] and adjusted for sex, age and BMI. Annotation of genomic location of differentially methylated sites was achieved with R package ChIPseeker [28]. Correlation analysis between methylation % and gene expression (log_2_) was conducted using the rcorr() function from Hmisc R package.

## Results

### Clinical characteristics

Full participant characteristics are displayed in **Table 1** and have been previously reported [15]. For contextualization of results, differences in phenotypic differences are detailed below. Active had a lower body mass (kg) in comparison to Obese (*P* < 0.05) and T2D (*P* < 0.01) which was due to lower fat mass (kg) (*P* < 0.001; **Table 1**). There were no body composition differences between T2D and Obese. By design, T2D had significantly reduced insulin sensitivity (M-value) in comparison to both Obese (*P* < 0.05) and Active (*P* < 0.001; **Table 1**). There were no differences in M-value between Active and Obese (**Table 1**). Aerobic capacity measured by VO2peak (L/min) was greater in Active compared to Obese and T2D (*P* < 0.001) with no differences between Obese and T2D (**Table 1**). This relationship persists when VO2peak is normalized to body mass (kg) or lean body mass (kg) (**Table 1**).

### Mitochondrial Content

Citrate synthase activity was not significantly different between Obese and T2D but was significantly greater in Active compared to T2D (*P* < 0.01; **Figure 1A**). mtDNA copy number was not significantly different between Obese and T2D but was greater in Active compared to both Obese and T2D (*P* < 0.001; **Figure 1B**).

**Figure 1.**
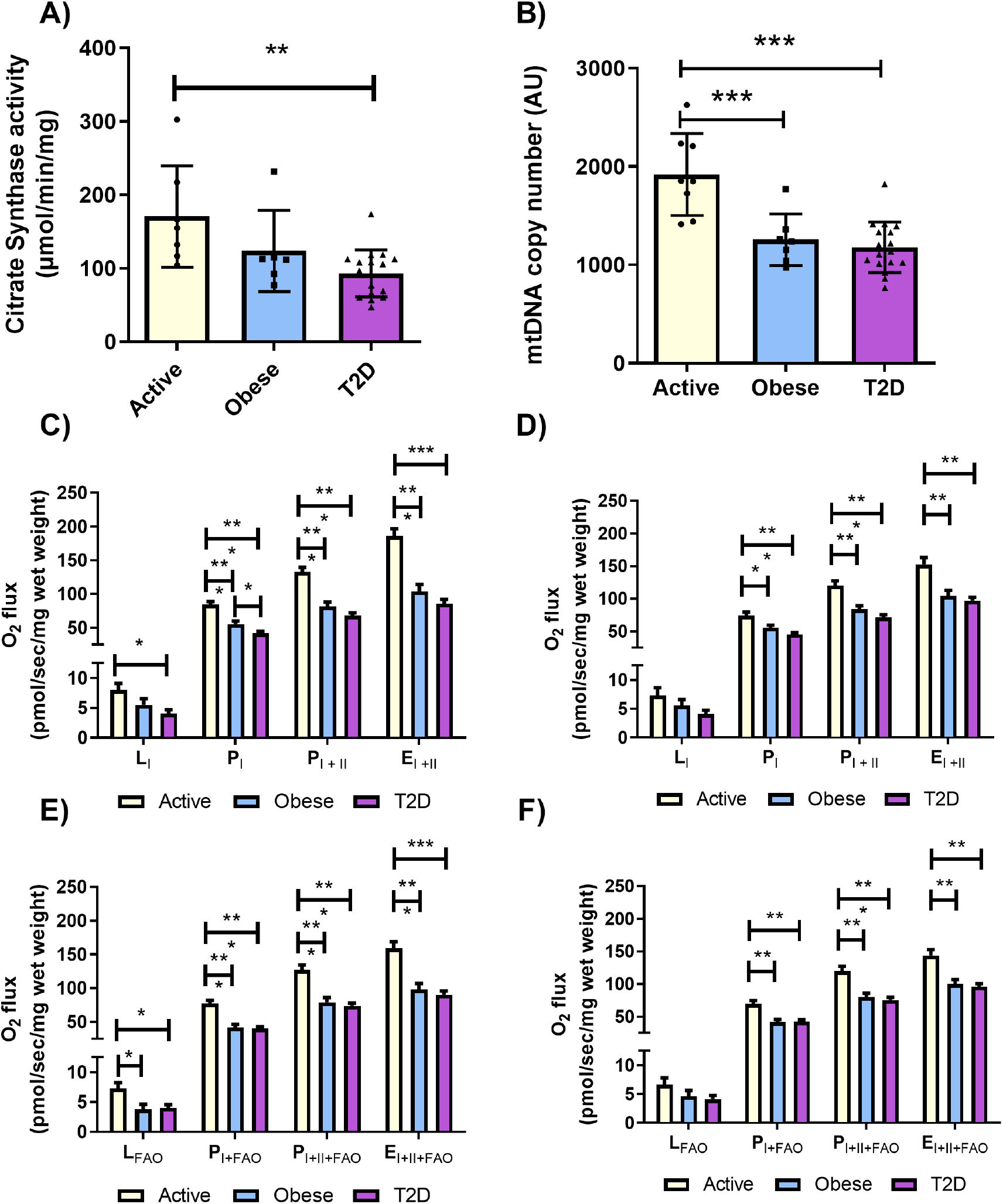
Differences in skeletal muscle mitochondrial content and *ex vivo* mitochondrial respiration adjusted for mitochondrial content between Active, Obese and T2D. Citrate synthase activity in skeletal muscle tissue (A). mtDNA copy number in skeletal muscle tissue (B). Data analyzed with ANOVA model with Tukey post hoc adjustment. Data are presented as mean ± SEM overlayed with individual values. Mitochondrial respiration quantified by high resolution respirometry in permeabilized skeletal muscle fibers during a glycolytic protocol, consisting of leak (L_I_), complex I supported OXPHOS (P_I_), complex I+II supported OXPHOS (P_I+II_), maximal electron transfer system capacity (E_I+II_) adjusted for citrate synthase activity (C) and mtDNA (D). Mitochondrial respiration during a fatty-acid oxidative protocol, consisting of leak (L_FAO_), complex I supported OXPHOS (P_I+FAO_), complex I+II supported OXPHOS (P_I+II+FAO_), maximal electron transfer capacity (E_I+II+FAO_) adjusted for citrate synthase activity (E) and mtDNA (F). An ANCOVA was used to analyze data with CS activity or mtDNA as a covariate followed by post-hoc comparisons. Data are adjusted means ± SE. *** *P* < 0.001, ** *P* < 0.01 * *P* < 0.05.

### *Ex vivo* skeletal muscle mitochondrial capacity

We previously showed that Active had greater O2 flux compared to Obese and T2D during respiratory states; P_I_, P_I+FAO_, P_I+II_, P_I+II+FAO_, E_I+II_, E_I+II+FAO_ [15]. There were no differences between Obese and T2D for any of these respiratory states. The differences between Active and Obese/T2D *ex vivo* glycolytic- and fatty-acid-supported mitochondrial capacities remained when respiration values were normalized to citrate synthase activity and mtDNA copy number (**Figure 1C-F**).

### Supercomplex formation

CIV-containing SCs were significantly greater in Active (n = 6) compared to Obese (n =4; *P* < 0.01) and T2D (n =15; *P* < 0.001; **Figure 2A-B**). CI- and CIII-containing SCs were higher in Active compared to T2D (*P* < 0.01; **Figure 2A-B).** There was also a trend (*P*=0.075) for CIII-containing SC being higher in Obese compared to T2D (**Figure 2B).** Unadjusted SC are shown in **supplementary figure 1**.

**Figure 2.**
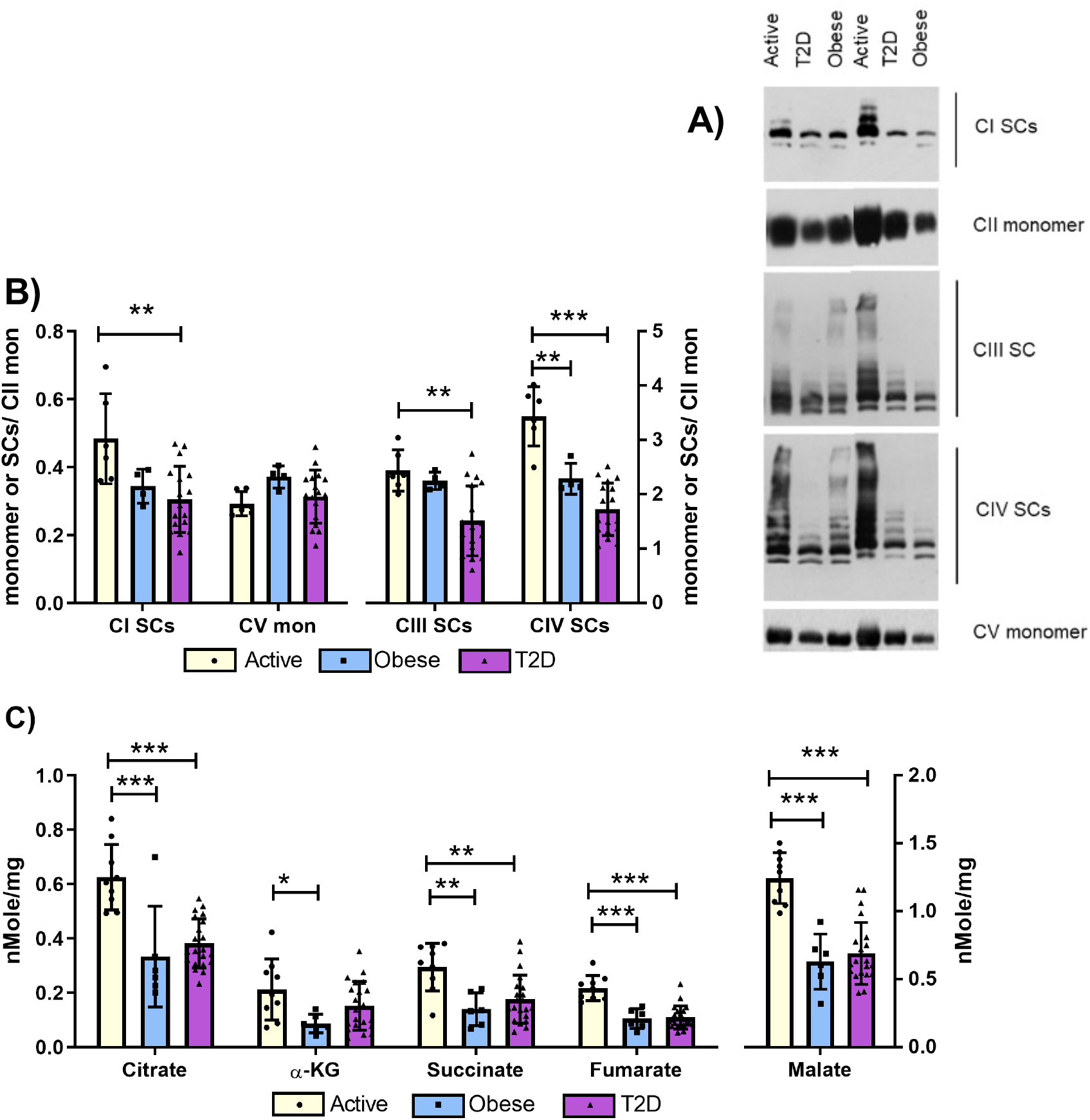
Differences in skeletal muscle supercomplex assembly and TCA intermediates between Active, Obese and T2D. Mitochondrial Supercomplex and monomers measured by BN-PAGE and quantified with antibodies; anti-NADH dehydrogenase [ubiquinone] 1 alpha subcomplex subunit 9, mitochondrial (NDUFA9; complex I), anti-flavoprotein (complex II), anti-ubiquinol-cytochrome c reductase core protein II (UQCRC2; complex III), anti-complex IV subunit I (complex IV) and anti-ATP synthase subunit alpha, mitochondrial (ATP5a; complex V) (A). Expression of the mitochondrial supercomplexes and monomers relative to complex II monomer (B). Quantification of the TCA cycle intermediates using target metabolomics (C). Data analyzed with ANOVA model with Tukey post hoc adjustment. Data are presented as mean ± SEM overlayed with individual values. *** *P* < 0.001, ** *P* < 0.01, * *P* < 0.05.

### TCAi

Citrate, succinate, fumarate and malate were all significantly greater in Active (n = 9) compared to Obese (n = 6; *P* < 0.001) and T2D (n = 19; *P* < 0.001; **Figure 2C**). α-ketoglutarate was significantly greater in Active compared to Obese only (*P* < 0.05; **Figure 2C**). There were no differences in any of the TCAi between Obese and T2D (**Figure 2C**). Apart from α-Ketoglutarate, these differences remained after the data were adjusted for citrate synthase activity (**Supplementary Figure 2A**). Differences also remained after the data were adjusted for mtDNA copy number with the exceptions of α-Ketoglutarate and succinate (**Supplementary Figure 2B**).

### Transcriptomic Analyses

Principal component analyses of top ranked contributing genes in the SkM showed clear separation in Active compared to T2D and Obese (**Figure 3A**), with considerable overlap between T2D and Obese. This separation in skeletal SkM was reflected by the number of differentially expressed genes (DEGs; *P* < 0.05) being much higher in T2D vs Active (489 downregulated, 2156 upregulated) and Obese vs Active (457 downregulated, 682 upregulated) than T2D vs Obese (276 downregulated, 593 upregulated; **Supplementary Tables 1-3**). Over-representation analysis showed Active had an upregulation of genes related to respiration/mitochondrial capacity and mitochondrial structure compared to both Obese and T2D (**Figure 3B, Supplementary Table 4**). In contrast, T2D had upregulation of genes related to protein degradation and catabolic processes compared to Active (**Figure 3B, Supplementary Table 4**); whereas, Obese had an upregulation of genes related to cell matrix and junction assembly compared to Active (**Figure 3B, Supplementary Table 4**). When comparing T2D against Obese there were no significant GO terms downregulated in T2D; whereas, protein targeting and localization were upregulated in T2D (**Figure 3B, Supplementary Table 4**)

**Figure 3.**
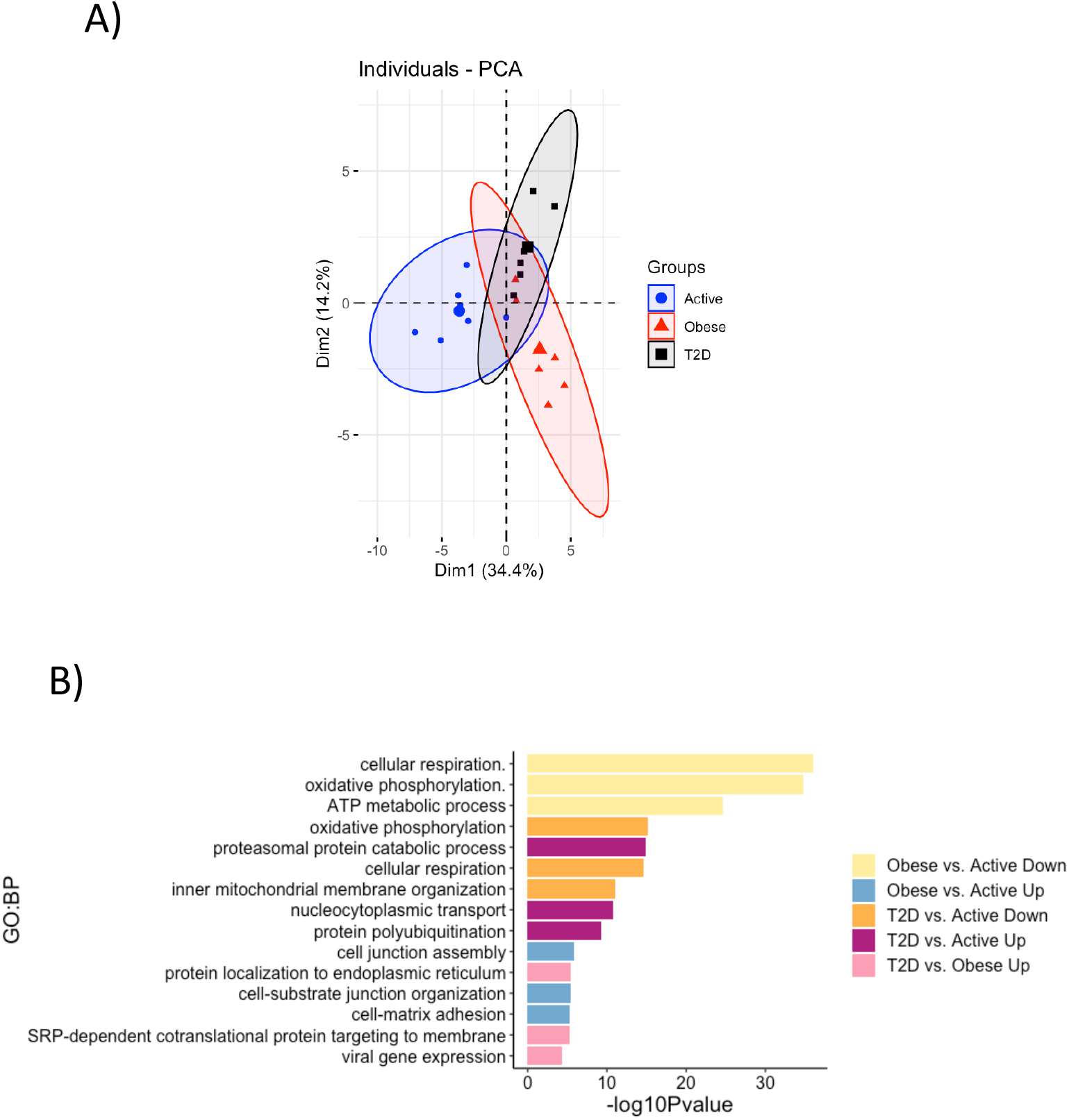
Transcriptomic differences between Active, Obese and T2D skeletal muscle. Principal component analysis of top ranked contributing genes in skeletal muscle (A) Selected gene ontology biological processes from over-representation analysis (B). N = 6 from each group.

### MitoCarta Analysis

We used the MitoCarta3.0 list [29] to identify which transcriptional aspects of mitochondrial capacity, structure and processes differ between Active, Obese and T2D. Out of a total 1136 MitoCarta genes, 169 were downregulated in Obese compared to Active and 143 were downregulated in T2D compared to Active (**Supplementary Table 5**). Downregulated genes in Obese and T2D were related to oxidative phosphorylation (OXPHOS) subunits and assembly factors, mitochondrial translation, metals and cofactors, lipid and carbohydrate metabolism and TCA cycle (**Figure 4**). A subset of MitoCarta genes (n=65) were upregulated in T2D compared to Active, which were largely related to autophagy, protein homeostasis and nucleotide metabolism (**Figure 4).** Eighty Mitocarta genes were upregulated in T2D compared to Obese which were related to aspects of various metabolic processes including, lipid metabolism, amino acid metabolism, FA oxidation, nucleotide metabolism and in some instances OXPHOS subunits and assembly factors (**Figure 4).** Obese had very few genes upregulated in comparison to Active (10 genes) and T2D (4 genes; **Supplementary Table 5**).

**Figure 4.**
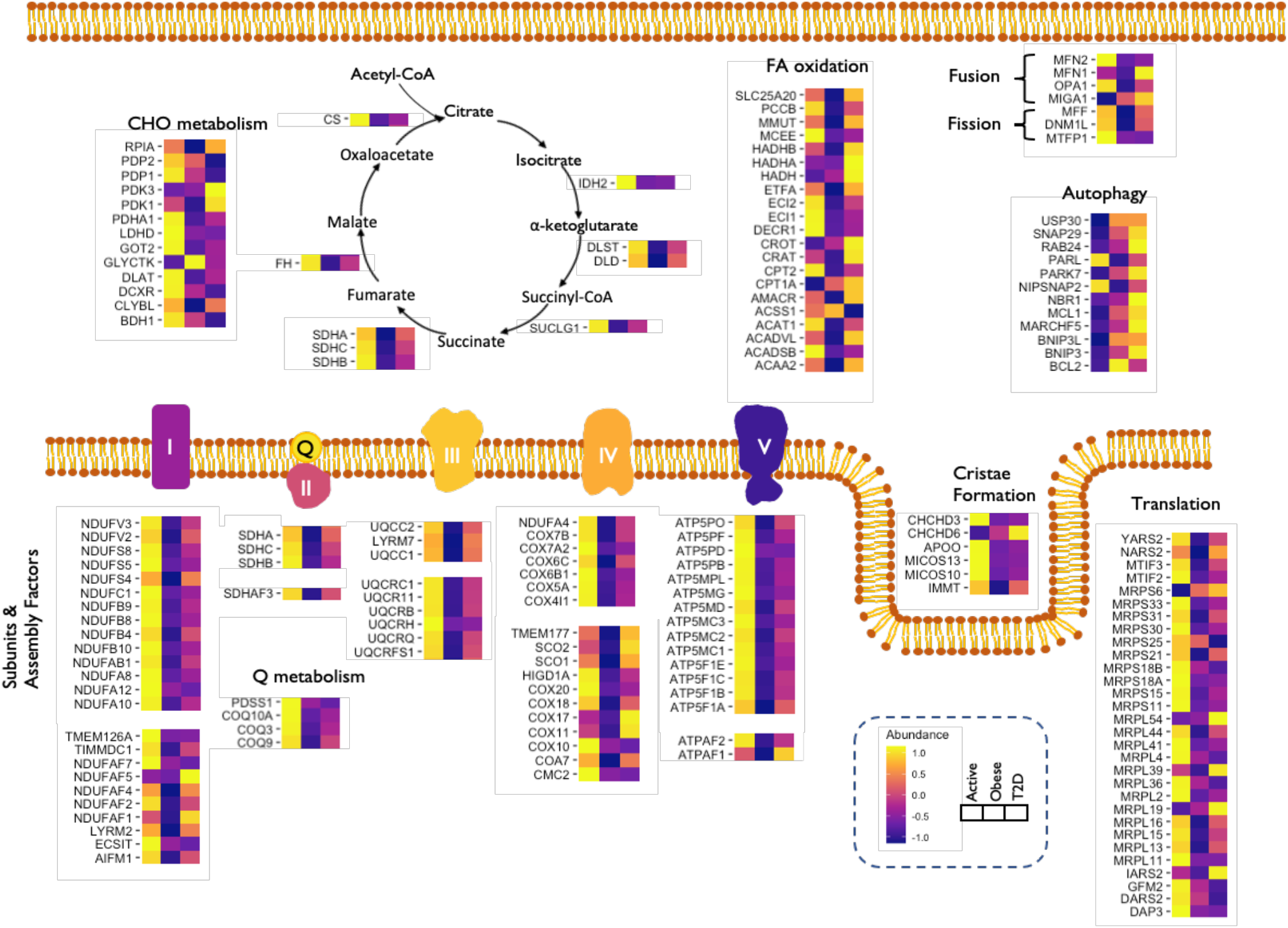
Heatmaps of MitoCarta DEGs between Active, Obese and T2D skeletal muscle. Data is presented as standardized average (n = 6 for each group) scores for each group.

### DNA Methylation Analyses

We determined how many MitoCarta DEGs contained differentially methylated CpG sites (DMS) for each comparison (**Figure 5A, Supplementary Tables 6**). Between 25% to 70% of MitoCarta DEGs had DMS (**Figure 5A**), with the highest absolute amount of DMS pertaining to downregulated genes in Obese compared to Active. We next ran correlation analyses to determine which DMS located at MitoCarta DEGs were correlated to their gene expressions for each group comparison (**Supplementary Table 7**). While each group comparison had similar percentages (25-35%) of DMS located at MitoCarta DEGs significantly correlated to gene expressions (**Supplementary Figure 3**), only 2% of DMS located at MitoCarta DEGs upregulated in T2D compared to Obese were correlated to gene expression (**Figure 5B**). There were only 2 DMS located at downregulated MitoCarta DEGS in T2D compared to Obese and neither correlated to gene expression.

**Figure 5.**
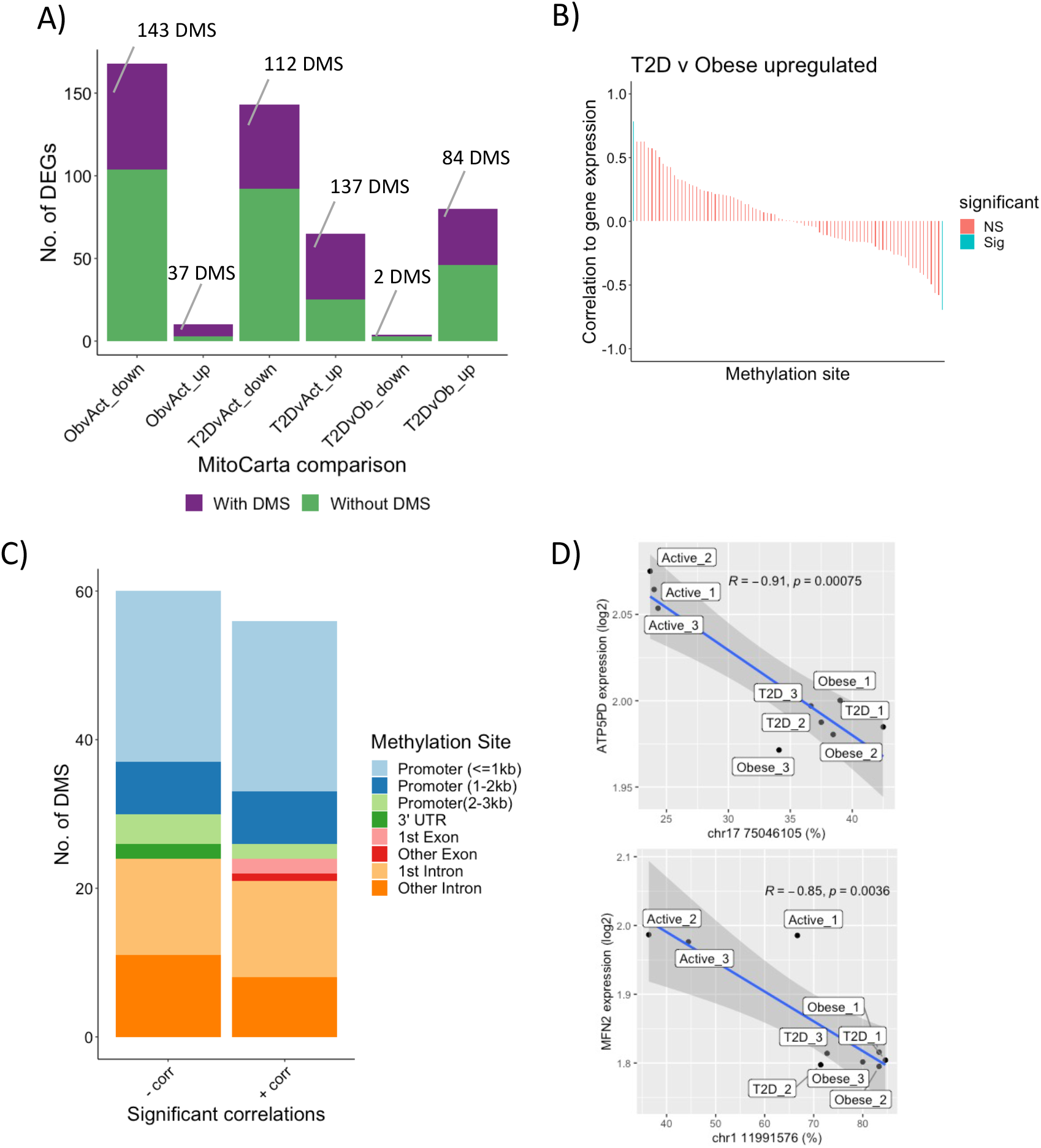
Differences in DNA methylation between Active, Obese and T2D in relation to differentially expressed MitoCarta genes. The number of Mitocarta differentially expressed genes (DEGs) for each comparison that have differentially methylated sites (DMS) and the number of DMS (A). Waterfall plot showing the correlation of DMS to MitoCarta DEGs and their significance (*P* < 0.05) (B). When correlations are split into comparisons T2D v Obese upregulated only showed 2 significant correlations (C). Genomic location of DMS that are correlated to differentially expressed MitoCarta genes (D). A total of 9 participants (n = 3 for each group) was used for analysis of DNA methylation data.

Independent of directionality of correlations to gene expressions, the majority of the DMS (79%) were located in either the promoter region or at the first intron (**Figure 5C**). There were a number of DMS negatively correlated with the expressions of genes encoding OXPHOS subunits and assembly factors which generally showed lower gene expressions and higher CpG methylation levels in Obese and T2D compared to Active (**Figure 5D**). Mitochondrial fusion gene *MFN2* had two promoter regions that were inversely correlated with gene expressions (one displayed in **Figure 5D**) and displayed lower gene expression and higher CpG methylation in Obese and T2D.

## Discussion

Insulin resistance and blunted mitochondrial capacity in SkM are often synonymous; however, this association remains controversial with previous research reporting conflicting results. Employing a comprehensive multi-factorial analysis of SkM mitochondrial capacity, we demonstrate that obese individuals with and without T2D have comparable mitochondrial capacities underscored by similarities in mitochondrial content, *ex vivo* respiration adjusted for mitochondrial content, supercomplex assembly and levels of TCA cycle intermediates. This lack of differences extends to the *in vivo* level where we previously demonstrated comparable PCr recovery rates between Obese and T2D [15]. A critical aspect of our study is that the Obese and T2D cohorts had similar levels of confounding factors such as BMI, age and aerobic capacity, which are known to impact mitochondrial capacity. Compared to sedentary individuals with obesity with and without T2D, we demonstrate that lean Active individuals with enhanced aerobic capacity have numerous aspects of superior SkM mitochondrial capacity quantified by mitochondrial content, *ex vivo* respiration adjusted for mitochondrial content, supercomplex assembly and levels of TCA cycle intermediates. By adjusting *ex vivo* respiration by mitochondrial content, we highlight that differences in overall mitochondrial capacity are in part due to enhanced intrinsic capacity of the mitochondria. These findings are paralleled by a robust upregulation of genes encoding crucial aspects of mitochondrial capacity (i.e. OXPHOS subunits and assembly factors, mitochondrial fusion, lipid and carbohydrate metabolism and TCA cycle). Furthermore, genes regulating OXPHOS subunits and mitochondrial fusion displayed reduced DNA methylation in the promoter regions correlating to enhanced gene expression.

In agreement with previous research, we show that individuals with obesity with and without T2D have comparable SkM mitochondrial capacities when measured *in vivo* or *ex vivo* [11]. This contrasts with previous reports showing reduced SkM mitochondrial capacity in individuals with T2D compared with BMI- and age-matched normo-glycemic controls when measured *in vivo* [30–32] or *ex vivo* [14,30]. The discrepancies among these studies can be explained by aerobic capacity not being controlled for [14,32] or the cohort investigated being overweight – rather than obese – and having higher aerobic capacities compared to our cohorts [30]. As expected, lean Active individuals had greater SkM mitochondrial capacity and was likely due to them being physically active and having greater aerobic capacity [33]. Interestingly, *ex vivo* respiration remained higher in Active and comparable between Obese and T2D when the results were adjusted for mitochondrial content measured by two indices. This contrasts previous research [34] and indicates greater SkM respiration in the Active group is not entirely due to greater mitochondrial content.

Research has demonstrated that the multi-protein complexes of the ETS do not simply exist as monomeric units in the mitochondrial inner membrane but instead can form higher order SC structures that contain varying proportions of CI, CIII and CIV. The formations of such SCs permit greater electron flow through the ETS, thus maximizing capacity and efficiency for ATP production [35–37]. The Active group had greater SC formation which aligns with previous research showing increases in SC assembly following endurance exercise training in previously sedentary individuals [37]. Our findings, however, are contrary to a previous report demonstrating reduced SC formations in individuals with T2D compared to BMI-matched non-diabetic group [19]. This previous study however used a different SkM group (*rectus abdominus*), participants with a significantly higher BMI and participants undergoing a hypocaloric diet for three weeks prior to bariatric surgery. The greater SC formation in Active supports the idea that mitochondrial capacity can be improved without changes in mitochondrial content and may be due to improved stoichiometry of SC formation.

The TCA cycle is the final common pathway for oxidation of substrates and the major source for generation of reducing equivalents for oxidative phosphorylation. In parallel with the previous results Active had greater TCAi and there were no comparable differences in TCAi between Obese and T2D. Most of the differences were retained after adjustment for mitochondrial content suggesting individual mitochondria have greater TCAi in Active. Scientific investigations of TCAi in human SkM are few in number. Succinate increases in human SkM of sedentary individuals following endurance training [38], aligning with our Active group. In mouse models, TCAi are reduced in lipid-induced insulin-resistant SkM [4] and in SkM of genetically obese mice, but to a lesser extent in genetically diabetic mice [39]. Together suggesting that TCAi are reduced in sedentary individuals compared to Active individuals.

Previous work has suggested a downregulation of genes related with OXPHOS in SkM of individuals with T2D [40]; however, this has not been recapitulated in larger clinical cohorts [41,42]. In agreement, we show that global transcriptomic differences between Obese and T2D SkM are not indicative of mitochondrial-related processes but rather appear to be related with protein targeting. OXPHOS gene expressions have been previously correlated with VO2peak [40]; thus, our observed upregulation of genes related to mitochondrial capacity in the Active group compared to Obese and T2D are likely due to their enhanced aerobic capacity – although we cannot discount the impact of obesity in our findings.

When a targeted approach was used to detect differences in mitochondria-specific genes [MitoCarta 3.0 [29]], it illustrated that Active had specific upregulation of numerous genes related to OXPHOS subunits and assembly factors, mitochondrial fusion and fission, mitochondrial translation, lipid and carbohydrate metabolism in comparison to Obese and T2D. Mitochondrial morphology, which is integral to optimal function is maintained by tightly regulated fusion and fission processes. Fission divides mitochondria and acts as a quality control for damaged mitochondria to undergo mitophagy [43]. Fusion binds mitochondria, resulting in mitochondrial elongation and enhanced cristae cross-sectional area. This promotes greater OXPHOS protein distribution throughout the cristae and allows for greater capacity of energy production [44]. Endurance exercise training is known to enhance regulators of mitochondrial fusion, MFN2 and OPA1 [45,46]. The upregulation of the fusion regulators (*MFN2* and *OPA1*) in our Active group, concurrent with upregulated OXHPOS subunits and assembly factors indicates mitochondria morphology is optimally maintained and contributes to enhanced function and oxidative phosphorylation. Interestingly T2D had an upregulation of a subset of MitoCarta genes linked to OXPHOS subunits and assembly factors, FA oxidation and lipid metabolism in comparison to Obese; however, this did not translate to any improvements in functional mitochondrial capacity.

DNA methylation is an epigenetic modification of DNA that can impact the regulation of gene transcription in human SkM [47]. Only 2% of DMS located at MitoCarta DEGs in the T2D compared to the Obese comparison correlated with gene expression. This is in stark contrast to 25-35% of DMS located at MitoCarta DEGs significantly correlated to gene expression for the remaining group comparisons (i.e. T2D vs Active, Obese vs Active etc); thus, suggesting that differential DNA methylation is not contributing to the observed differences in transcriptional regulation of mitochondrial capacity between T2D and Obese. We found that independent of directionality of correlations to gene expressions, the majority of DMS (79%) were located in either the promoter region or at the first intron which is near the transcription start site (TSS) [48]. We observed increased gene expression with increased methylation of DNA in almost half the instances. The repressive role of promoter DNA methylation on gene transcription has long been established [49]. It is worth noting that transcription factors have varying degrees of sensitivity to CpG methylation with only 22% exhibiting decreased binding to their motifs with hypermethylation [50]. Therefore, it is plausible that the DMS positively correlated to gene expressions do not have reduced transcription factor binding and have increased transcription through other epigenetic modifications. A number of critical mitochondrial-related genes linked to OXPHOS subunits and mitochondrial fusion had a negative correlation between methylation and expression that were upregulated in the Active group. Together, this indicates that major components of mitochondrial capacity that are enhanced in the Active group are regulated at the level of DNA methylation.

The Obese group had normal HbA1c (%) (5.7%) and greater insulin-stimulated glucose uptake (M-Value) compared to the T2D group, which was comparable to the Active group. Therefore, our T2D group was insulin resistant compared to our BMI-, aerobic capacity- and age-similar controls. It is noteworthy that T2D diagnosis duration was on average 4.6 years, which along with the high HbA1c (%), places them at a reduced probability of remission [51,52] (i.e. they are at a ‘later stage’ of T2D progression). Therefore, SkM insulin resistance and reduced glycemic control precedes the development of mitochondrial dysfunction in this group of individuals with T2D.

By leveraging a comparison to an Active lean group, we were able to highlight molecular, stoichiometry and functional differences that contribute to superior SKM mitochondrial capacity. We hypothesize that this superior mitochondrial capacity is due to their enhanced aerobic capacity [33]. Future research should include a lean sedentary control group to assess if obesity contributes to impaired mitochondrial capacity rather than aerobic capacity. We aimed to recruit both sexes in all cohorts; however, we ended up with an uneven sex distribution, with the Active group comprised of only Males and the majority of Obese being females. Females tend to have superior mitochondrial function than males [53]. Despite this finding, our Obese group had similar mitochondrial capacity to T2D and the lowest expression of mitochondrial genes. We conclude the differences we observed are due to group differences and are not impacted by sex distribution.

In summary, our highly controlled, in-depth multi-factorial analysis dissociates SkM mitochondrial capacity from insulin resistance with robust support from *ex vivo* respiration, targeted metabolomics, SC assembly, global transcriptomics and DNA methylation. We further highlight that lean, active individuals have enhanced SkM mitochondrial capacity compared to sedentary obese individuals with and without T2D at the *ex vivo*, metabolomics and global transcriptomics levels that is linked in part to differential DNA methylation.

## Acknowledgments

We thank the study volunteers for their participation and the TRI clinical research staff for their contributions.

## Data availability

Raw and processed RNA-Seq and DNA methylation data sets generated and analyzed during the current study are available in the NCBI GEO repository (GSE196387). All other data generated and analyzed during the current study are available from the corresponding author upon reasonable request. No applicable resources were generated or analyzed during the current study.

## Authors’ relationships and Activities

The authors have nothing to disclose.

## Author Contributions

Conceptualization, KLW, LMS, SRS and MW; Methodology, HC, GP, MFP, DAP; Software KLW, GY and YS; Formal Analysis, KLW, MFP, GY, YS, FGDC, AD, FY, HC, GP, DAP, MEH, RBV; Investigation, KLW, MFP, YS, RY, GP, DAP, HC, SG; Resources, GP, DAP, MEH MFP; Data Curation; KLW, MFP, GY, YS, FGDC, FY, NK; Writing – Original Draft, KLW; Writing Review & Editing, GY, YS, RY, GP, AD, HC, DAP, MEH, MFP, SRS, MW, LMS, NK. Visualization, KLW, GY, GP; Supervision, MEH, SG, SRS, MW, LMS; Project Administration, MFP, LMS; Funding Acquisition, LMS.

## Abbreviations

DEGS: Differentially Expressed Genes
DMS: Differentially Methylated Sites
ETS: electron transport system
GO: gene ontology
OXHPOS: oxidative phosphorylation
PCA: Principal Component Analysis
PCr: Phosphocreatine
RRBS: Reduced representation bisulfite sequencing
SC: supercomplex
SkM: Skeletal Muscle
TCA: Tricarboxylic

---

**Supplementary Figure 1.**
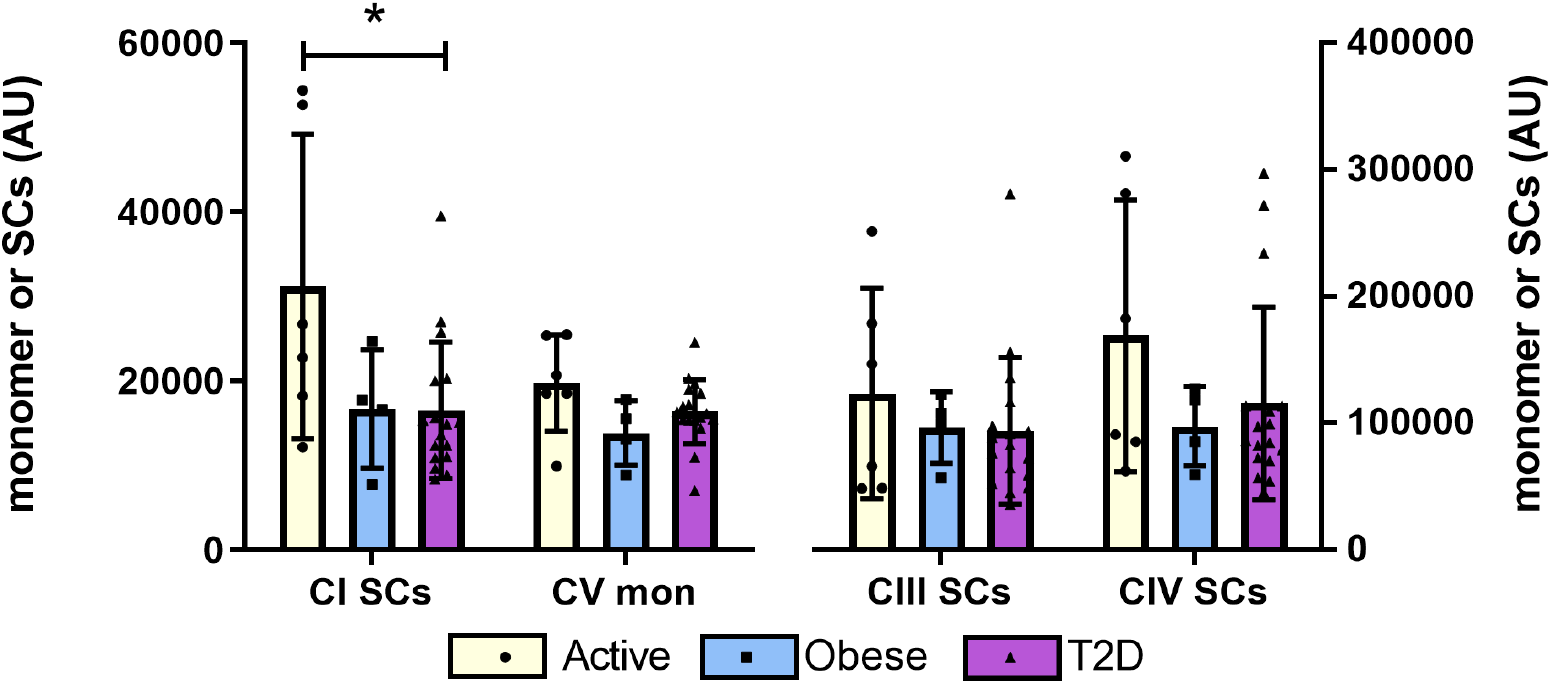
Unadjusted supercomplex assembly. Raw expression of the mitochondrial supercomplexes and monomers. Data are presented as mean ± SEM overlayed with individual values. * *P* < 0.05.

**Supplementary Figure 2.**
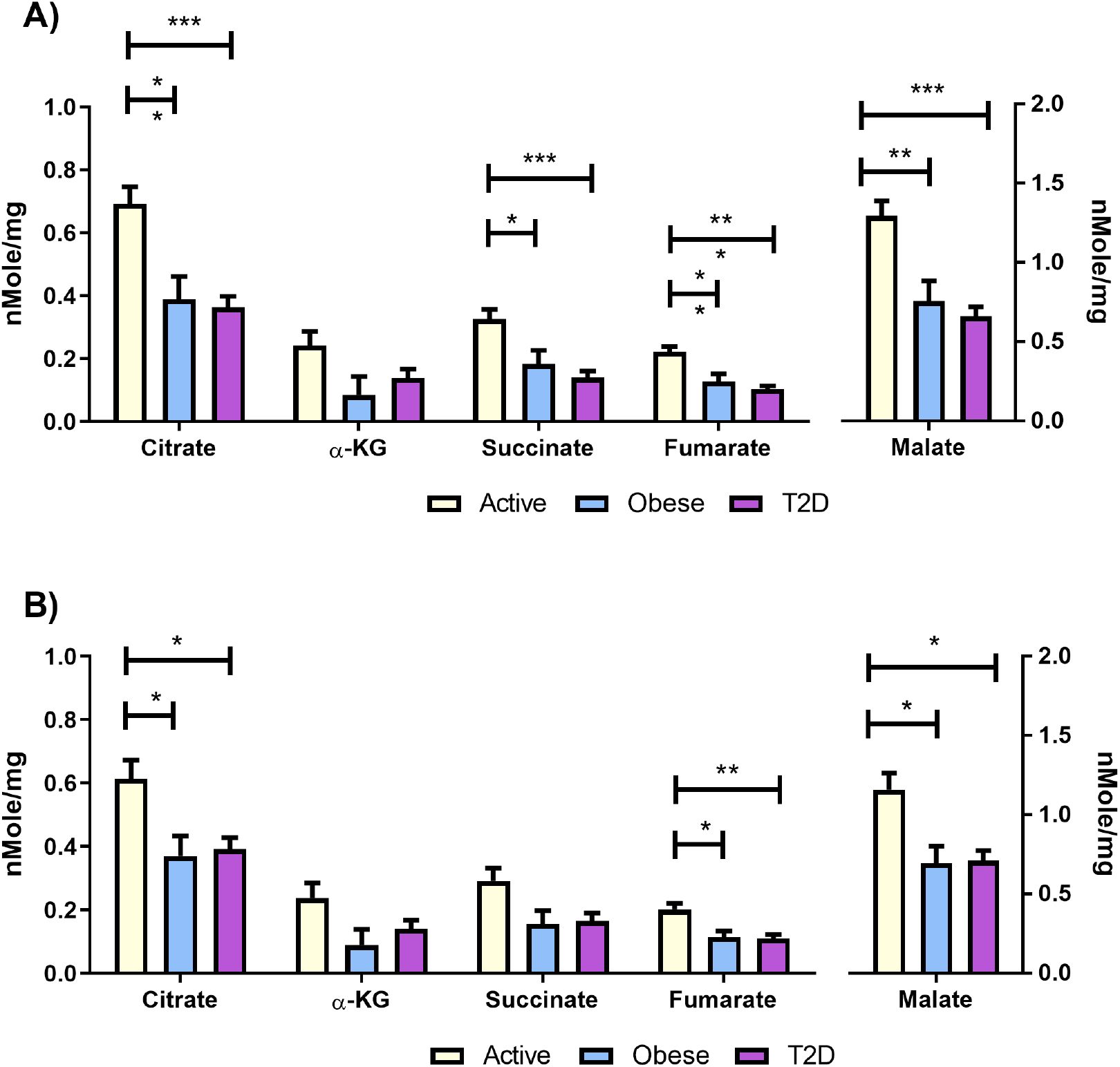
TCA cycle intermediates adjusted for mitochondrial content. An ANCOVA was just to analyze data with CS activity as a covariate followed by post-hoc comparisons (A). An ANCOVA was just to analyze data with mtDNA as a covariate followed by post-hoc comparisons (B). Data are adjusted means ± SEM. *** *P* < 0.001, ** *P* <0.01, * *P* <0.05.

**Supplementary Figure 3.**
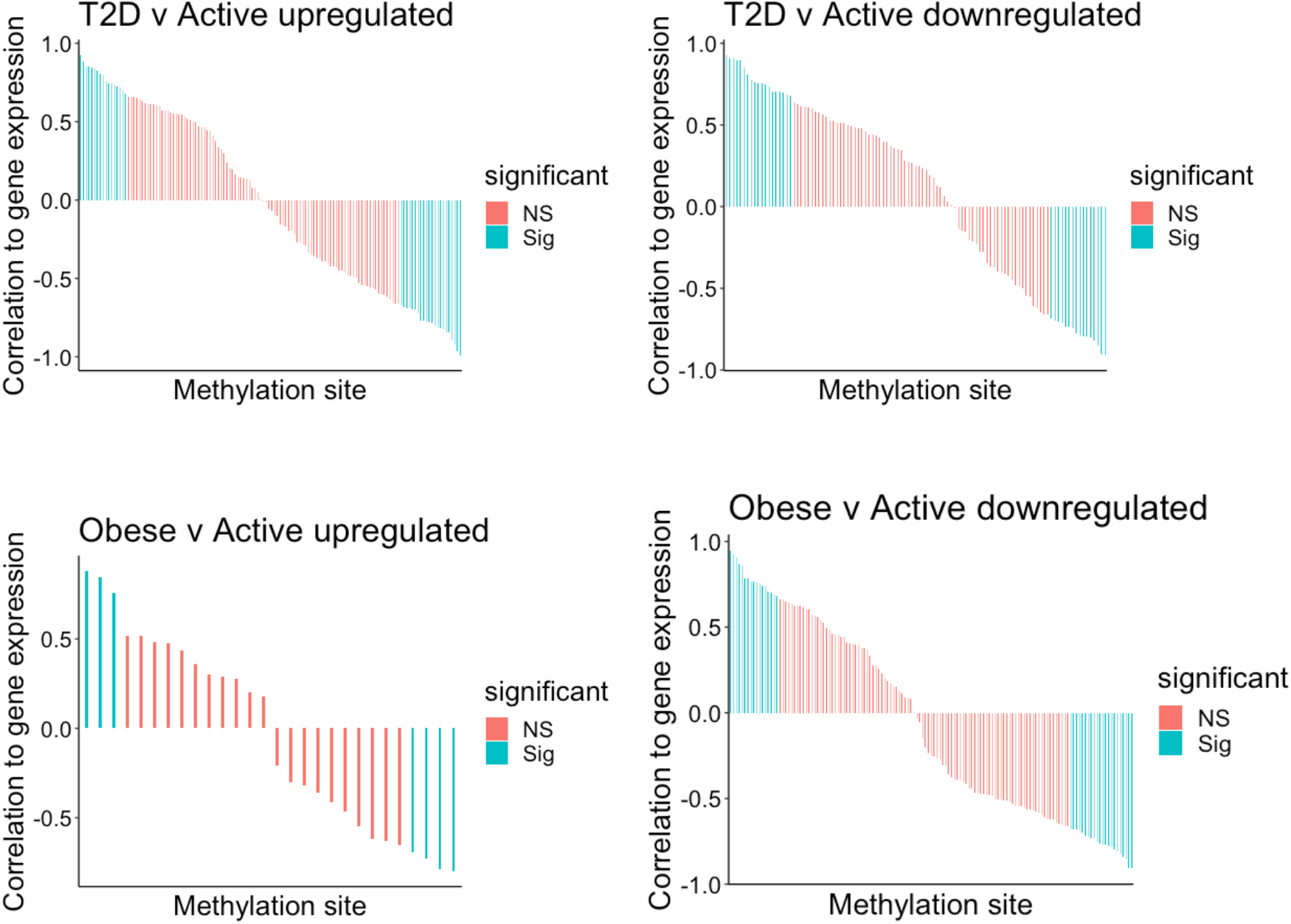
Waterfall plots showing the correlation of DMS to MitoCarta DEGs and their significance (*P* < 0.05) for each comparison. A total of 9 participants (n = 3 for each group) was used in the correlation analysis.

